# Estimating variance components in population scale family trees

**DOI:** 10.1101/256396

**Authors:** Tal Shor, Dan Geiger, Yaniv Erlich, Omer Weissbrod

## Abstract

The rapid digitization of genealogical and medical records enables the assembly of extremely large pedigree records spanning millions of individuals and trillions of pairs of relatives. Such pedigrees provide the opportunity to investigate the sociological and epidemiological history of human populations in scales much larger than previously possible. Linear mixed models (LMMs) are routinely used to analyze extremely large animal and plant pedigrees for the purposes of selective breeding. However, LMMs have not been previously applied to analyze population-scale human family trees. Here, we present **S**parse **C**holesky factor**I**zation LMM (Sci-LMM), a modeling framework for studying population-scale family trees that combines techniques from the animal and plant breeding literature and from human genetics literature. The proposed framework can construct a matrix of relationships between trillions of pairs of individuals and fit the corresponding LMM in several hours. We demonstrate the capabilities of Sci-LMM via simulation studies and by estimating the heritability of longevity and of reproductive fitness (quantified via number of children) in a large pedigree spanning millions of individuals and over five centuries of human history. Sci-LMM provides a unified framework for investigating the epidemiological history of human populations via genealogical records.

**Author Summary:** The advent of online genealogy services allows the assembly of population-scale family trees, spanning millions of individuals and centuries of human history. Such datasets enable answering genetic epidemiology questions on unprecedented scales. Here we present Sci-LMM, a pedigree analysis framework that combines techniques from animal and plant breeding research and from human genetics research for large-scale pedigree analysis. We apply Sci-LMM to analyze population-scale human genealogical records, spanning trillions of relationships. We have made both Sci-LMM and an anonymized dataset of millions of individuals freely available to download, making the analysis of population-scale human family trees widely accessible to the research community. Together, these resources allow researchers to investigate genetic and epidemiological questions on an unprecedented scale.

## Introduction

Genealogical records can reflect social and cultural structures, and record the flow of genetic material throughout history. In recent years, very large pedigree records have come into existence, owing to collaborative digitization of large genealogical records[1,2] and to digitization of large cohorts collected by healthcare providers, spanning up to millions of individuals[3–7]. Such population-scale pedigrees allow investigating the sociological and epidemiological history of human populations on a scale that is orders of magnitude larger than existing studies.

Traditional human pedigree studies collect a large number of independent families which are analyzed separately and then meta-analyzed. However, this approach is not suitable for population-scale pedigrees, because such pedigrees cannot be decomposed into mutually exclusive families[1]. Hence, the analysis of such pedigrees requires modeling complex covariance structures between trillions of pairs of individuals.

Pedigree studies often employ LMMs to decompose the phenotypic variation among individuals into variance components such as genetic effects and shared environment[8]. LMMs have been the statistical backbone of animal and plant breeding programs for almost six decades[9], and have been continuously developed over the years[10–21]. LMMs are routinely used nowadays to analyze pedigrees of millions of animals and plants[13,22], hundreds of thousands of which are often genotyped (e.g. [23,24]).

In recent years, LMMs and their extensions have also gained considerable popularity in human genetics studies for the purposes of estimating heritability[25–31] and genetic correlation[32–36], predicting phenotypes[37–40] and modeling sample relatedness[41–45]. Unlike classical animal and plant studies, human studies typically do not include pedigree data, but instead measure genetic relatedness via dense genotyping of single nucleotide polymorphisms (SNPs).

In recent years, animal and human studies used different techniques to scale LMMs to datasets with millions of individuals. Animal studies typically fit large-scale LMMs via restricted maximum likelihood (REML)[12], by exploiting the sparsity of pedigree data. Specifically, a pair of individuals with no known common ancestor can be regarded as having no genetic similarity. Consequently, these pairs induce a zero entry in the genetic similarity matrix. Such sparse matrices can be stored and analyzed efficiently with suitable numerical techniques[21,46,47].

Human genotyping studies do not give rise to sparse data structures. Instead, human studies have managed to scale LMMs to large datasets via two approaches. The first approach applies REML, either via supercomputers with thousands of CPUs and terabytes of memory[45], or by approximating the restricted likelihood gradient via Monte-Carlo techniques[27]. However, the latter technique is only suitable for specific types of covariance matrices whose decomposition is known beforehand.

The second approach to scale LMMs uses the method of moments rather than REML, by solving a set of second moment matching equations[48–52]. Such approaches have become increasingly popular recently[29–34,53–57] owing to their computational tractability and their compatibility with privacy-preserving summary statistics[58]. Although moment estimators are less statistically efficient compared to REML estimators, they have several advantages: the loss of efficiency has been found to often be small[55]; they are more robust to modeling violation because they make fewer distributional assumptions; and they are more flexible, which enables applying technique to limit confounding factors such as assortative mating (Methods). Moment estimators have also recently been explored in animal breeding studies[59–61] and were found to be faster than REML while providing similar accuracy, but they have not been widely adopted in animal studies to date.

Here we propose Sci-LMM, a statistical framework for analyzing population-size pedigrees that combines techniques from animal and human genetic studies. Sci-LMM uses sparse data structures as is common in animal studies, and supports both moment and REML estimators. The moment estimator is based on a common technique called Haseman-Elston (HE) regression[62,63] (Methods). Sci-LMM scales HE regression to population-sized pedigrees via sparse matrix tools[64]. The REML estimator combines a direct sparse REML solver [46] with Monte-Carlo gradient approximation[65]. Importantly, existing packages for pedigree-based REML[66–70] cannot handle the analyses performed in this paper because they require the inverse of the epistatic interactions matrix[46,71,72], which is extremely difficult to compute in large pedigrees[73]. Hence, Sci-LMM provides a comprehensive solution for LMM-based pedigree analysis.

To demonstrate the capabilities of Sci-LMM, we carry out an extensive analysis of simulated data with millions of individuals, which we complete within a few hours. We additionally estimate the heritability of longevity and of reproductive fitness (quantified via number of children), using a large cohort spanning millions of genealogical records and several centuries of human history. We estimate that both traits have a substantial heritable component, with an estimated 22.1% heritability for longevity and 34.4% for reproductive fitness. Sci-LMM enables analysis of large pedigree records that was not previously possible.

## Results

### Overview of Proposed Framework

#### Sci-LMM allows decomposing a range of factors contributing to phenotypic variance

Sci-LMM is a flexible framework to analyze population scale pedigrees with various types of covariates and variance components. Sci-LMM assumes that phenotypes follow a multivariate normal distribution, whose mean is affected by fixed effects associated with covariates such as age and sex, and whose covariance is affected by variance components associated with dependency structures such as familial relations or geographical proximity. The aim is to identify the contribution of these fixed effects and variance components and to decipher their relative contribution. For example, when modeling both geographical proximity and familiar relationships, the magnitudes of the variance components can indicate whether the studied trait is more strongly transmitted via inheritance or via environmental factors. Sci-LMM estimates these parameters using either REML or HE regression (Methods).

Following the classical genetic model originally proposed by Fisher[74] and developed by Cockerham[71] and Kempthorne[72], Sci-LMM supports the efficient computation of an additive IBD-based genetic component, a dominancy, and an epistasic covariance matrix (Methods). These matrices can be constructed rapidly via dynamic programming[75–77].

In addition to covariance matrices, Sci-LMM also includes adjustments using principal components (PCs) with fixed effects, which can be computed efficiently using sparse matrix routines[78]. These adjustments can capture major linear sources of variation in a dataset that may not be captured well via the entire covariance matrix, such as founder effects. However, we note that unlike PCs computed from genetic data, PCs computed from IBD are not guaranteed to capture population structure[79].

### Simulation Studies

To evaluate the capabilities of Sci-LMM, we generated large synthetic pedigrees spanning 20 to 40 generations and various family structures, under a wide variety of settings. The pedigrees included 50,000-2,000,000 individuals, amounting to trillions of pairs of relatives. A subset of the individuals in each generation consists of children of individuals from either the previous generation, or from two generations in the past. To simulate patterns observed in real datasets, the simulations also included consanguinity, half-siblings, and individuals with less than two recorded parents (Supplementary Material).

Next, we used the pedigree structure to compute the additive, epistasis and dominance matrices. We then generated a normally distributed phenotype, using ten binary covariates and a covariance matrix given by a weighted combination of the above matrices. Unless otherwise stated, the sparsity factor (the fraction of non-zero entries in each matrix) was 0.001. We generated ten different datasets for each combination of sample size and sparsity factor that we investigated (Supplementary Material).

In all settings, Sci-LMM yielded empirically unbiased estimates of the variance components, using both REML and HE regression. As expected, estimation accuracy increased with sample size, though the estimators became slightly less accurate when increasing the number of variance components, (Figure 1a-c). Specifically, the root mean square error (RMSE) was < 0.03 for all methods under all settings with more than 250,000 individuals, indicating <3% average error (because the phenotype was standardized to have unit variance).

**Figure 1:**
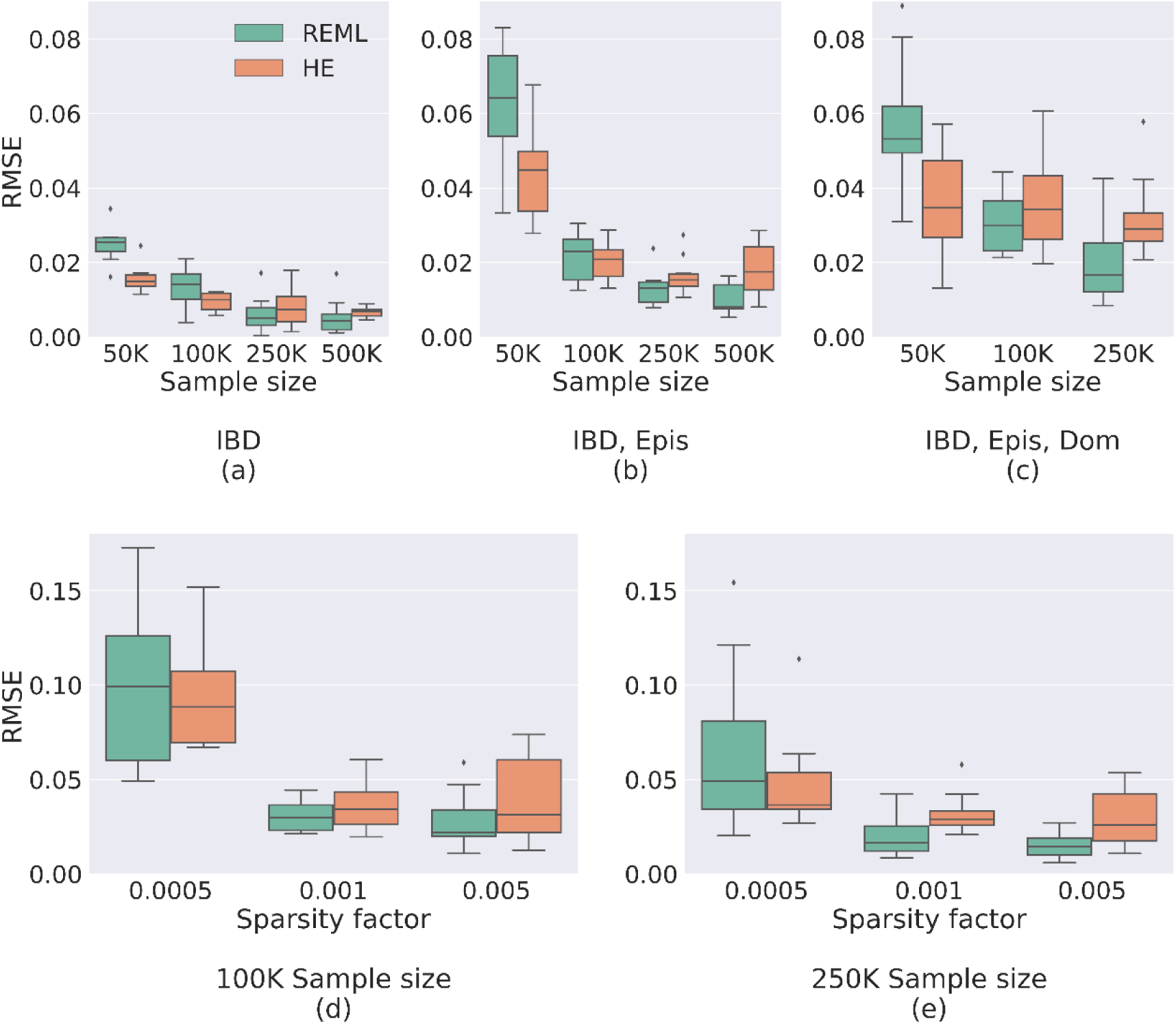
Evaluating the estimation accuracy of Sci-LMM. (**a-c**) Box plots comparing REML and HE estimation accuracy (RMSE) across simulated datasets (each box represents 10 experiments), under varying sample sizes, using **(a)** only IBD, **(b)** IBD and epistasis, or **(c)** IBD, epistasis and dominance variance components. HE is more accurate than REML for smaller sample sizes, but REML outperforms HE as the sample size increases. Results for analyses with three matrices and 500,000 individuals are omitted due to excessive required computational time. (**d-e**) Comparing REML and HE estimation accuracy when using IBD, epistasis and dominance matrices under various sparsity factors (the fraction of non-zero matrix entries) with either **(d)** 100,000 individuals, or (**e**) 250,000 individuals. The estimation accuracy of both REML and HE increases with the number of non-zero entries, for both REML and HE.

A comparison of the REML and the HE results shows that that HE was slightly more accurate in the presence of <100,000 individuals (Figure 1a-c), and REML was slightly more accurate otherwise. These results possibly indicate that REML convergence is difficult in the presence of sparse covariance matrices with limited sample sizes. We also found that estimation accuracy was anti-correlated with relatedness sparsity, indicating that the estimators efficiently exploit the information found in non-zero covariance entries (Figure 1d-e).

#### Runtime

Next, we evaluated the matrix construction time. While Sci-LMM supports several variance components, the rate limiting factor is the IBD matrix construction. We found that the Sci-LMM run-time scales linearly with the number of non-zero entries in the matrix (Figure 2a-b). For example, Sci-LMM takes less than 4 hours to construct an IBD matrix with 5 × 10^11^ pairs of possible relatives and a sparsity factor of 0.001.

**Figure 2:**
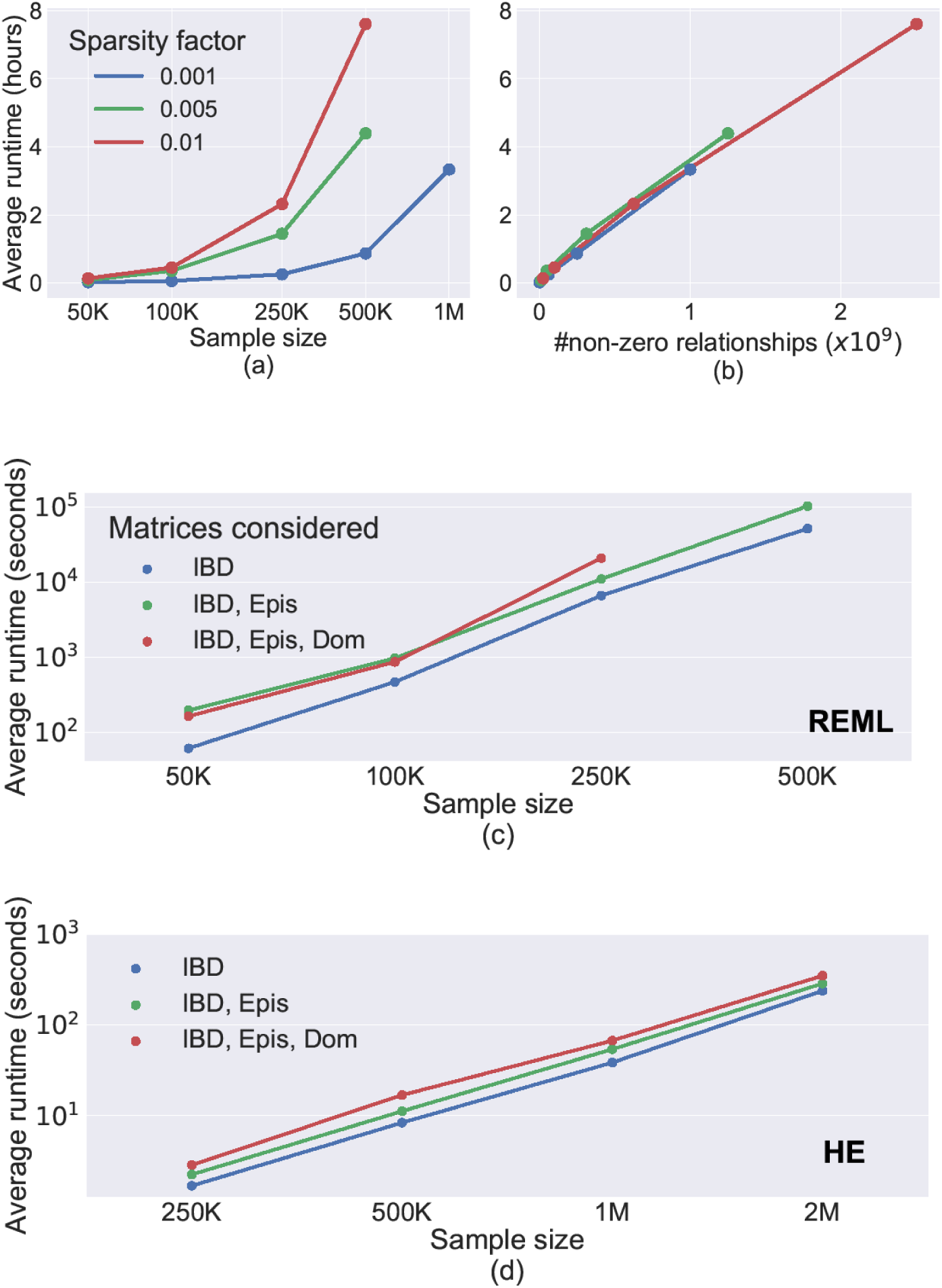
Analysis of Sci-LMM computation time. **(a)** Computation time required to compute an IBD matrix from pedigree data under different sparsity factors as a function of sample size. **(b)** Computation time required to compute an IBD matrix from pedigree data as a function of the number of nonzero relationships, demonstrating a linear relationship. The maximal number of evaluated non-zero relationships increases with the sparsity cutoff, because we only generated matrices with up to a million individuals. **(c)** Variance component estimation time (using REML), as a function of sample size, when using different combinations of covariance matrices. Epis – Epistasis; Dom - dominance **(d)** same as (c), but for HE regression instead of REML estimation. Here we evaluated datasets with up to 2 million individuals that were not investigated in (c), owing to technical limitations of the sparse matrix factorization routines used in our REML implementation.

Next, we investigated REML and HE estimation runtime. REML estimation for samples with 500,000 individuals (representing 250 million covariance entries) required less than 24 hours (Figure 2c), whereas HE estimation required 16 seconds (Figure 2d). Overall, our results demonstrate that the Sci-LMM framework is scalable to extremely large pedigrees.

Finally, we tried comparing Sci-LMM with pedigree-based REML software packages[66–69]. We could not invoke any of these packages with an epistatic covariance matrix because they require its inverse, which is not sufficiently sparse[73]. We verified this by trying to invert the matrix analytically via the software package nadiv[80] and numerically via sparse matrix libraries[81] and via the matrix inversion facilities of the software package WOMBAT[68], all of which ran out of memory on a 256GB machine. We additionally tried running the analysis via the *fitNullModel* function of the GENESIS package[82] and the *lmekin* function of the coxme package[83], both of which can work with sparse covariance matrices. However, both packages could not finish the analysis in 4 days, presumably because they do not use the approximate gradient approximation techniques of Sci-LMM.

When excluding the analysis to only an additive covariance matrix, WOMBAT was much faster than Sci-LMM, completing an analysis of 250,000 individuals in several minutes (Supplementary Table 1). This is because REML estimation with only an additive matrix can be extremely efficient when combining the analytical form of the inverse matrix[75–77] with a mixed model equations solver[46].

#### Estimating the heritability of longevity and reproductive fitness

We used Sci-LMM to estimate the heritability of longevity and reproductive fitness, based on large-scale pedigree records obtained from the Geni genealogical website[1] (see Web Resources). An initial description of the longevity analysis was reported in [1], but here we substantially refine and extend this analysis. We applied stringent quality control to minimize deaths due to non-natural reasons such as wars or natural disasters, by excluding pairs of individuals who died within 10 days of each other or within periods with significantly elevated death rates[1]. This filtering yielded approximately 441,000 individuals with birth and death dates. We first computed the IBD, dominance and epistasis matrices of these individuals, and then estimated the heritability of longevity using these matrices.

The corresponding IBD matrix contained over 3 billion nonzero entries. It included the 441,000 core individuals and their informative ancestors, yielding a total of 1.6 million individuals. The submatrix consisting of only the core individuals included 251 million non-zero IBD coefficients (yielding a sparsity factor of ∼0.001, in correspondence with the simulation studies). The dataset included 9.7 million pairs of individuals with a kinship coefficient corresponding to a >=20-degree relationship (Figure 3a). Sci-LMM constructed this matrix in 10 hours.

**Figure 3:**
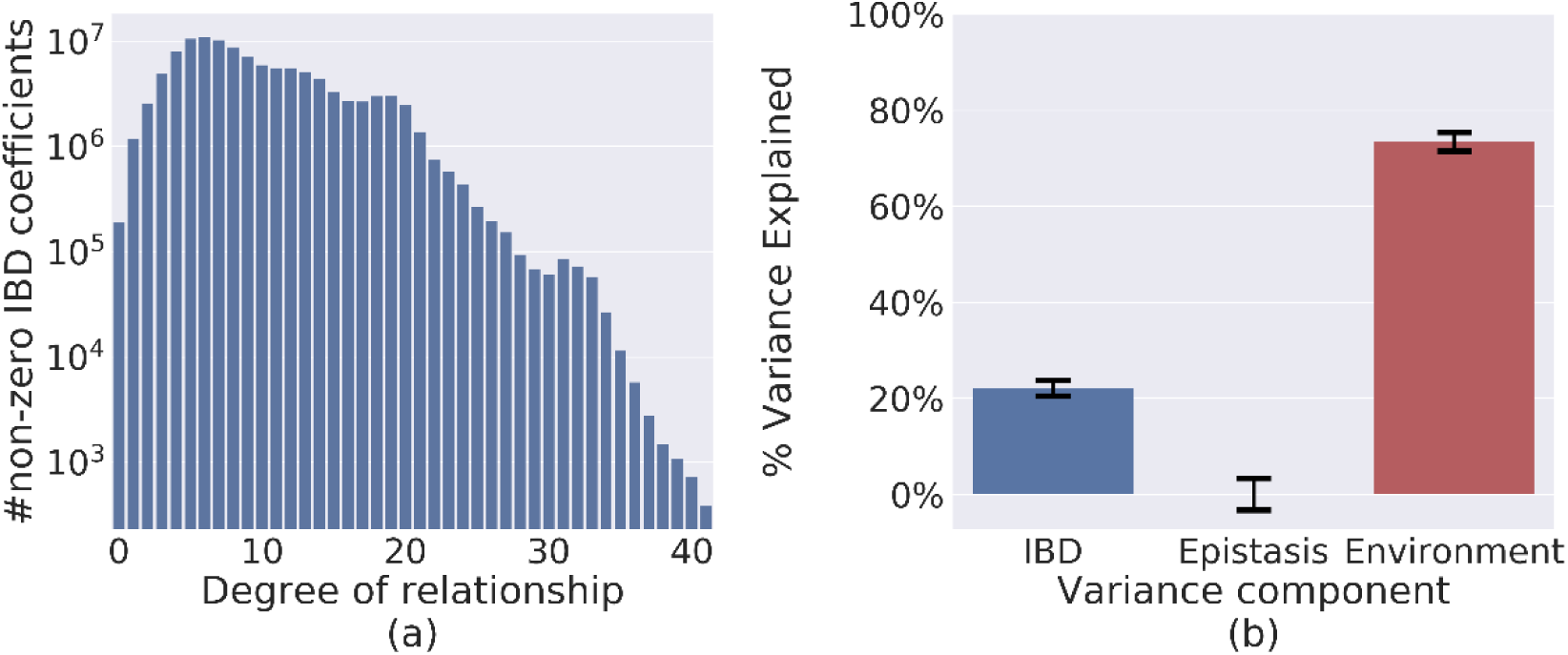
Results of analysis of a real pedigree with 441,000 individuals. **(a)** A histogram of genetic similarity across 441,000 individuals, using only the closest relationship between every pair of individuals. The dataset includes approximately 9.7 million pairs of individuals whose least common ancestor lived at least 10 generations earlier. **(b)** The estimated fraction of longevity variance attributed to different variance components (and their 95% CI).

Next, we used Sci-LMM to estimate the heritability of longevity with covariates encoding gender, year of birth (yob), yob^2^, yob^3^, and the top 10 principal components of the IBD matrix, and with covariance matrices encoding IBD and pairwise epistasis (Methods). A dominance matrix was not included because the analysis included a relatively small number of parent-child relationships, rendering this matrix almost equivalent to the identity matrix (Supplementary Material). The REML estimates were: IBD: 22.1% (s.e. 0.8%); pair-wise epistasis: 0.001% (s.e. 1.7%); environmental effects: 74.4% (s.e. 1.0%) (Figure 3b). The HE estimate for IBD was 24.3% (s.e. 0.5%) when excluding the epistatic interactions matrix (HE results with epistatic interactions were inconclusive due to large standard errors). We also tried to limit confounding due to shared environmental effects[2,84] by excluding ∼399,000 pairs (∼0.15%) of individuals with a shared household (spouses or parent-child pairs) from the analysis without excluding the individuals themselves, using HE (Methods). This led to a heritability estimate of 26.3% (s.e. 0.9%), indicating that shared household effects are unlikely to up-bias our estimates. Overall, our results suggest that the heritability of longevity is upper bounded by ∼26%. However, the true heritability may be lower since our estimates may be confounded by other environmental factors[2,84] (see Discussion).

We also estimated the heritability of reproductive fitness, quantified by number of children. To limit confounding due to non-genetic factors, we applied stringent filtering of individuals. We removed individuals with less than two children records, individuals with a shared household (spouses and children) and individuals who are first-or second-degree relatives with another individual in the data set. The filtered dataset included ∼45,000 individuals. We used the same covariates as before, applied a Box-cox transformation to induce normality for number of children, and excluded epistatic interactions from this reduced dataset, because they led to large standard errors. The REML and HE estimated heritability of reproductive fitness were 28.4% (s.e. 0.5%) and 34.4% (s.e. 1.2%), respectively. These results indicate a substantial genetic component for reproductive fitness, in line with previous work[85]. As before, we caution that this estimate may be upper-biased due to confounding by non-genetic factors[2,84] (see Discussion).

Finally, we performed a series of experiments to examine the robustness of our analysis to potential confounding factors. These included considering only individuals born after 1800, excluding first-and second-degree relatives from the analysis, excluding individuals that share or likely shared a household in the past, and various combinations thereof. The results indicate that our analysis is not upper biased due to inclusion of these potential confounding factors (Supplementary Table 2).

## Discussion

We have described a statistical framework for analysis of large pedigree records spanning millions of individuals. Our framework includes methodologies for constructing large sparse matrices given raw pedigree data, and methodologies for LMM analysis with random effects described by these matrices. Taken together, the proposed solution enables an end-to-end analysis of population scale human family trees.

In this work we focused on partitioning phenotypic variance into additive genetics, epistasis and dominance. However, the LMM framework is flexible and can be extended in various directions. For example, sparse LMMs are often used to model transmissible phenomena [86–91], which enables combining pedigree-based and geography-based covariance structures. Both Sci-LMM and the data studied in this paper are freely available for download, which makes the analysis of population-scale human family trees widely accessible to the research community. Combined, these resources allow researchers to investigate genetic and epidemiological questions on unprecedented scales.

We evaluated two methods for variance components estimation: REML and HE regression. REML is more accurate than HE and provides a likelihood-based solution, which can be used for model comparison and hypothesis testing. HE estimates are less accurate but can be more robust to modeling violation. Importantly, HE regression can mitigate confounding due to environmental factors by zeroing selected entries in the covariance matrices, which may be especially suitable for studying human genealogical records (Methods). Hence, the two methods are complementary in terms of their strengths and weaknesses. In practice, we found that it is difficult to scale REML to datasets with more than 500,000 individuals with a sparsity factor of 0.001. Our recommendation is to use REML when it is feasible and all model assumptions hold, and HE regression otherwise. We note that REML estimation can be substantially faster when not fitting epistatic interactions by using a mixed model equations approach [46], which is implemented in several software packages[66–69].

Our work demonstrates the technical feasibility of studying population scale human family trees. However, the analysis of human genealogical records is challenging due to imperfect data and the difficulty of controlling for confounding factors. Potential issues include non-paternity, cryptic relatedness, missing or false genealogical records, genetic nurture[92,93], environmental bias[84,94], assortative mating[2] and correlation between additive and epistatic effects[17,18]. As such, our estimates should be considered as a first order approximation, and our heritability estimates are likely upper biased due to confounding. We expect that several recently proposed techniques to address these issues (e.g. [2,93,94]) could be integrated into the Sci-LMM framework in the future.

In this work we focused on analyzing large pedigree records with no measured genotypes. In recent years, the advent of biobank-sized datasets allows analyzing population-scale genotyped cohorts. The two study types are complementary because biobanks cannot be used to investigate longevity, traits with a late age of onset, or epidemiological and sociological questions on historical scales. We anticipate that cohorts combining both types of data will become increasingly common. For example, we and other online genealogy platforms allow users to upload their genetic information and link it with their genealogical profile. Such combined datasets have been extensively explored in the animal breeding literature[19,21,95–99]. However, privacy and logistical concerns limit public access to human genetic data, necessitating methods based on summary statistics[58]. Thus, approaches for analysis of such combined datasets will combine state of the art techniques from the animal breeding and the GWAS literature, and remain a potential avenue for future work.

## Material and Methods

In the following paragraphs, we present the basic elements of each one of our algorithms. For brevity, the text describes the most important aspects necessary to understand our approach. The Supplementary Material presents the full technical details.

### Linear Mixed Models

Consider a sample of *n* individuals with observed phenotypes *y*_1_, …, *y*_*n*_, and covariates vectors ***C***_1_, …, ***C***_*n*_, and consider a set of *n* × *n* covariance matrices *M*^1^, …, *M*^*d*^, where 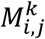 encodes the covariance between the phenotypes of individuals *i* and *j* according to the *k*^th^ covariance structure, up to a scaling constant. Under LMMs, the vector ***y*** = [*y*_1_, …, *y*_*n*_]^*T*^ follows a multivariate normal distribution:

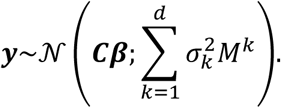

Here, ***C*** = [***C***_1_, …, ***C***_*n*_]^*T*^ is an *n* × *c* matrix of covariates (including an intercept), ***β*** is a *c* × 1 vector of fixed effects, and 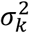 is the k^th^ variance component. Typically, one of the covariance matrices is the identity matrix, which encodes all sources of variance that is not captured by the other matrices.

### Computing an Identical by Descent Matrix

The IBD (identity by descent) kinship coefficient of two individuals, denoted as *a*_*ij*_, is the probability that a randomly selected allele in an autosomal locus was originated from the same chromosome of a shared ancestor between individuals *i* and *j* [100,101], and is given by:

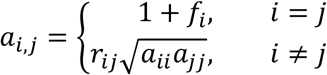

Here, *f*_*i*_ is the inbreeding coefficient, defined as half of the IBD coefficient of the parents of individual *i* [100], and *r*_*ij*_ is the coefficient of relationships, defined as:

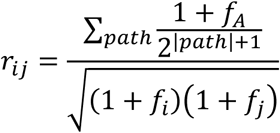

The quantities in the above equation are defined as follows: *A* is a least common ancestor of individual *i* and *j* in the pedigree graph (a graph where every node is an individual connected to her parents and children); the summation is performed over every path connecting individuals *i, j* in the pedigree graph, culminating at some ancestor A, such that the path does not contain the same individual twice; and |path| is the number of individuals on a path.

To efficiently compute the IBD matrix we first construct the matrices ***L*** and ***H*** of its decomposition ***A*** = ***LHL***^***T***^, where ***L*** is a lower triangular matrix such that ***L***_***ij***_ contains the fraction of genome shared between individuals *i* and her ancestor *j*, ***H*** is diagonal, and the matrices are ordered such that ancestors precede their descendants [75] (Figure 4a-c). The matrices ***L*** and ***H*** can be computed efficiently via dynamic programming [77] with sparse matrix routines [81].

**Figure 4:**
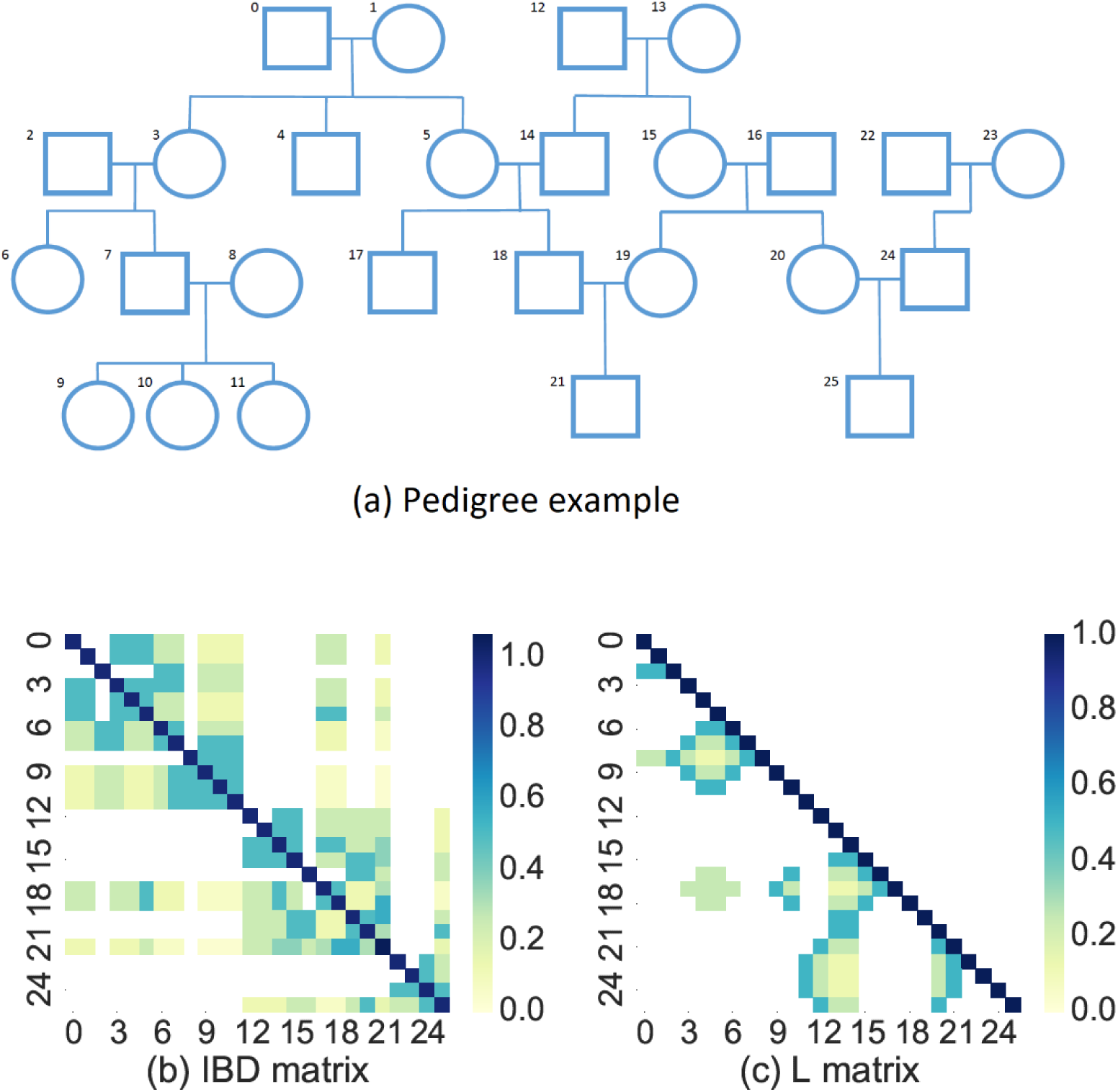
A demonstration of the Sci-LMM IBD matrix construction algorithm. (**a**) An example pedigree with 26 individuals. (**b**) A heat-map representing the IBD matrix, where zero elements are white to emphasize sparsity. (**c**) A heat-map representing the lower Cholesky factorization of the IBD matrix (i.e. the matrix *L* in the factorization ***A*** = ***LDL***^*T*^, where ***A*** is the IBD matrix). The value of entry *i, j* is the expected fraction of the genome that is shared between individual *i* and her ancestor *j*.

### Computing a Dominance Relationship Matrix

Dominancy represents the genetic variance due to co-ancestry of two alleles, and can be approximated by 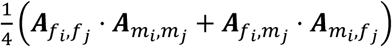, where *A*_*k,l*_ is the IBD coefficient of individuals *k, l*, and *f*_*k*_, *m*_*k*_ are the parents of individual k [10,102]. A necessary condition for nonzero dominancy entry is a nonzero IBD relationship, which enables rapid computation of the dominance matrix.

### Computing an Epistatic Relationship Matrix

Epistatic covariance encodes the assumption that variants interact multiplicatively to affect a given phenotype, and is proportional to the exponent of the corresponding IBD coefficient, i.e., (*A*_*k,l*_)^2^ for two-loci epistasis, (*A*_*k,l*_)^3^ for three-loci epistasis and so on [72]. Therefore, an epistatic covariance matrix can be computed in a straightforward manner given the IBD matrix.

### Pruning of uninformative individuals

Population scale pedigree data typically presents heterogeneity of the completeness of records. However, individuals with missing data may still be required for IBD computation. For example, consider a pedigree of two siblings with phenotypic data, and two parents and one uncle without phenotypic data. The parents are important for the IBD computation of the siblings, but the uncle is non-informative.

Sci-LMM applies pedigree-pruning techniques to remove non-informative individuals, similarly to other REML packages for pedigree analysis[67–70]. Briefly, we defined required individuals as individuals who have phenotypic and explanatory variables data, or individuals who appear in a lineage path connecting two individuals with such data with one of their least common ancestors. This algorithm reduces the matrix construction time by several hours.

### Haseman-Elston Regression

HE regression estimates variance components via the method of moments, using:

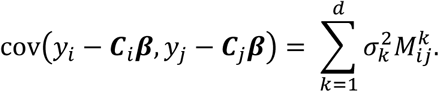

This allows fitting the variance components 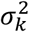 via least square minimization, which can be solved via sparse matrix routines. Typically, the fixed effects ***β*** are first estimated without considering the covariance matrices (which yields a consistent estimator under mild regularity conditions[103]), and are then incorporated into the moment estimator above. The sampling variance can be efficiently approximated via Month-Carlo approximations, similar to REML gradients (Supplementary Material).

HE regression provides a simple technique for excluding specific pairs of individuals (e.g. spouses) from the analysis without excluding the individuals themselves. This can be useful when trying to limit confounding due to factors such as assortative mating. Excluded pairs can be omitted from the analysis by zeroing the covariance matrix entries of corresponding pairs. Importantly, this technique cannot be used in REML, since the resulting covariance matrices may not be positive definite. We note that another potential approach to capture environmental risk factors is by including shared effects with a suitable incidence matrix[10], but this approach requires additional assumptions and has not been used here.

### Restricted Maximum Likelihood Estimator (REML)

The LMM restricted log likelihood (after discarding constant terms) is given by:

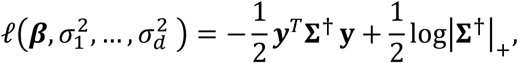

where **∑**^†^ is the Moore-Penrose pseudo-inverse of the covariance matrix 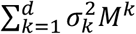 after projecting out the covariates (Supplementary Material), and |**∑**^†^|_+_ is the product of its non-zero eigenvalues[104]. This expression, as well as its partial derivative according to each variance component, can be computed efficiently via sparse matrix routines. Exact gradient estimation is more computationally challenging, but can be approximated efficiently via Monte Carlo techniques described in[27,65]. Since REML is sensitive to initial values, we obtain initial estimates via HE regression.

### Computing environment

All experiments were conducted using a Linux workstation with a 24-cores 2GHz Xeon E5 processor and 256Gb of RAM.

## Supporting information

## Acknowledgements

We thank Regev Schweiger, Elior Rahmani, Eran Halperin and Saharon Rosset for fruitful discussion. This study was supported by a generous gift from Andria and Paul Heafy (Y.E) and the Burroughs Wellcome Fund Career Awards at the Scientific Interface (Y.E.).

## Supporting Information Legends

**Supplementary Material**: Detailed derivations and algorithms for the methodologies described in the main text.

