## Supplementary material for "Estimating variance components in population scale family trees"

### Supplementary Material: Estimating variance components in population scale family trees

**4 Real Data Analysis** **7**

**5 Haseman Elston Estimator** **8**

**6 Restricted Maximum Likelihood Estimator** **9**

**8 Estimating Standard Errors of Variance Component Estimates** **14**

### 1 Supplementary Tables

Supplementary Table 1: Comparison of Sci-LMM and WOMBAT [16] runtime and memory requirements, when using simulations with only an additive IBD matrix. Both tools used only 1 CPU thread, and WOMBAT was executed with the --meuwissen flag. The estimated values were essentially the same for both tools in all cases. WOMBAT and Sci-LMM REML both crashed in the presence of pedigrees with ≥500K individuals.

| Sample size | Method | Time (minutes) | Peak memory (Megabytes) |
|---|---|---|---|
| 10000 | Sci-LMM (IBD computation) | 0.23 | 111 |
| | Sci-LMM (HE) | 0.02 | 68 |
| | Sci-LMM (REML) | 0.02 | 101 |
| | WOMBAT | 0.03 | 232 |
| 100000 | Sci-LMM (IBD computation) | 4.0 | 306 |
| | Sci-LMM (HE) | 1.07 | 631 |
| | Sci-LMM (REML) | 2.3 | 1462 |
| | WOMBAT | 1.3 | 232 |
| 250000 | Sci-LMM (IBD computation) | 14.0 | 1032 |
| | Sci-LMM (HE) | 5.3 | 2201 |
| | Sci-LMM (REML) | 150.0 | 12443 |
| | WOMBAT | 12.7 | 471 |
| 500000 | Sci-LMM (IBD computation) | 27.0 | 3621 |
| | Sci-LMM (HE) | 12.75 | 8904 |
| | Sci-LMM (REML) | - | - |
| | WOMBAT | - | - |

Supplementary Table 2: Sci-LMM heritability of longevity estimates under different analysis approaches. The epistasis estimates were very close to zero in all cases and are omitted for clarity.

| Individuals to exclude | sample size | IBD (HE) | IBD (REML) |
|---|---|---|---|
| born before 1800 | 283073 | 0.23 (0.006) | 0.23 (0.004) |
| in same household | 276011 | 0.29 (0.006) | 0.28 (0.005) |
| 1st or 2nd degree relatives | 110237 | 0.53 (0.040) | 0.51 (0.040) |
| In same household or $\leq$ 2nd degree relatives | 106469 | 0.539 (0.045) | 0.51 (0.044) |

### 2 Data Simulations

#### 2.1 Generating pedigrees

We generated pedigrees mimicking real marriage and child-bearing patterns in the United States, partially based on publications by the United States Census Bureau [18, 19].

We iteratively generated generations of individuals, where the first generation included two individuals, and the number of individuals in each successive generation increases by 40% (approximately the same ratio as in the GENI dataset [10]), until obtaining the desired sample size. Each generation included 50% females and 50% males.

In each generation we generated households, where every household includes either one individual or two individuals with different genders, and every individual can belong to zero, one or multiple households. The number of households in each generation was 62.5% of the number of individuals in that generation. 68% of the households included pairs of individuals, and the rest included a single individual.

Every individual in every generation (except for the top one) was born to parents from a randomly selected household from the previous generation (for 80% of individuals) or from two generations in the past (for the remaining 20% of individuals).

Finally, we omitted randomly selected edges until obtaining the desired sparsity factor, up to a 10% error.

#### 2.2 Generating covariance matrices

After generating a pedigree, we created corresponding IBD, dominance and epistasis matrices, as described in the main text.

#### 2.3 Generating a phenotypes

The final stage in each simulation consisted of generating a phenotype for every individual. We did this as follows. We first generated a variance components $\sigma_k^2$ for every covariance matrix $\mathbf{M}^k$ (including the identity matrix) from $U[0,1]$, and scaled them such that they sum to 1.0. We also generated a matrix $\mathbf{C}$ of 10 covariates, 5 of which were binary (with 50% probability of being 0), and 5 were sampled from a standard normal distribution. We also generated a vector $\boldsymbol{\beta}$ of covariate effects from $\mathcal{N}(0, 1000/n)$, where $n$ is the sample size.

Next, we sampled a phenotype vector $\mathbf{y}$ from its distribution:

$$\mathbf{y} \sim \mathcal{N}(\boldsymbol{C\beta},\ \sum_{k=1}^{d} \sigma_k^2 \mathbf{M}^k), \tag{1}$$

where $d$ is the number of matrices used.

To do this, we computed the Cholesky factorization of the aggregated covariance matrix, $\sum_{k=1}^{d} \sigma_k^2 \mathbf{M}^k = \mathbf{L}\mathbf{L}^T$ (where $L$ is a lower-diagonal matrix) via the CHOLMOD package, and then computed $\mathbf{y} = \mathbf{L}\mathbf{v} + \boldsymbol{C\beta}$, where $\mathbf{v}$ is a vector of iid normally distributed variables.

#### 2.4 Simulation parameters

The parameters differentiating the various experiments are the following:

1. Cohort size (50K, 100K, 250K, 500K, 1M or 2M).

2. Sparsity factor, which is the fraction of nonzero elements in the IBD matrix (efficiently computed using results of Lemma 1). The evaluated value were 0.0005, 0.001, or 0.005. The pedigrees were generated to an accuracy of up to 10% of the desired sparsity factor by removing random edges.
3. The subset of matrices used. All experiments used an IBD matrix, and a subset of the experiments additionally used a pair-wise epistasis [11] and a dominance inherritance matrix [8].

We generated 10 different datasets for every unique combination of settings, with the exception of matrices with 2M individuals, for which we generated a single pedigree with ten different phenotype vectors due to runtime considerations.

### 3 Computation of Covariance Matrices

#### 3.1 IBD matrix

We computed an IBD matrix that encodes IBD coefficients between all pairs of individuals with year of birth, year of death and geographical location in two stages. First, we pruned the dataset to retain only informative individuals. Afterwards, we computed the IBD matrix via an efficient dynamic programming algorithm. We now describe these two stages in detail.

##### 3.1.1 Removing non-informative individuals

The Familinx dataset [10] includes $N_0 = 43M$ individuals. Of these, only $N = 441K$ were selected by Kaplanis *et al.* as eligible individuals (i.e., individuals who passed various filtering criteria, such as not likely to have died due to non-natural causes, who have records of year of birth and year of death, etc.). However, a subset of the non-eligible individuals is still required for IBD computation purposes. For example, an individual with no year of birth, that is a parent of two eligible individuals, is still required for encoding the information that these two individuals are siblings.

We therefore distinguish between *eligible* individuals, which are the individuals selected by Kaplanis *et al.* and *informative individuals*, who are either (1) eligible, or (2) non-eligible but are required for IBD computation purposes.

The first stage of our IBD computation procedure consists of removing uninformative individuals. By using the definition of IBD (defined in the main text), the list of informative individuals consists of (1) eligible individuals; and (2) individuals who appear in a path connecting two eligible individuals with their least common ancestors.

We performed the pruning in three steps. First, we sorted individuals such that every individual precedes her offspring, using the networkx package [6].

Second, we removed non-eligible individuals who are not ancestors of any eligible individual (Supplementary Figures 1a, 1b). To perform this pruning efficiently, we created a matrix $AM$ such that (1) $AM_{ij} > 0$ only if there is a path of parent-child links connecting individuals $i$ and $j$; and (2) $i > j$ only if individual $j$ is not an offspring of

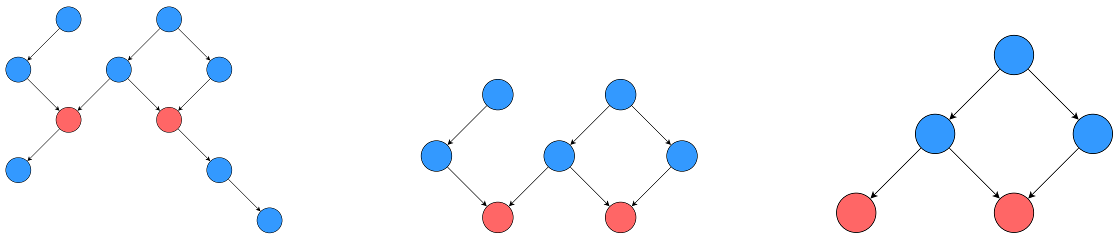

(a) Original pedigree

(b) Removing uninformative offspring

(c) Removing uninformative ancestors

Supplementary Figure 1: Stages of removal of uninformative individuals. Nodes represent individuals, and edges represent parent-child relations. Only red individuals have full information records (e.g. year of birth, year of death, etc.)

individual $i$. By denoting $rel$ as a binary adjacency matrix of parent-child pairs with the same ordering, we can compute the desired matrix $AM$ as follows:

$$AM = \sum_{i=1}^{oa} rel^i, \tag{2}$$

where $oa$ represents the longest path between an individual and her ancestor. This expression exploits the fact that for an adjacency matrix $M$, the matrix $M^d$ (for some integer $d > 0$) is a matrix of distances, such that $M_{i,j}$ is the number of paths of length $d$ between individuals $i$ and $j$. The $AM$ matrix can be efficiently computed with compressed row matrices [1].

Given the matrix $AM$, we can trivially remove all non-eligible individuals that have no eligible offspring.

In the next step, we removed non-eligible individuals who are not in any path between two eligible individuals passing through a common ancestor (Figure 1c). For this, we computed the matrix $CAM = (AM + I) \times (AM + I)^T$ and used the results of Lemma 1 to efficiently find all such individuals.

**Lemma 1.** *For every pair of individuals $i$ and $j$, $CAM[i, j] > 0$ if and only if $i$ and $j$ share common ancestors, where $CAM$ is defined as $(AM + I) \times (AM + I)^T$*

*Proof.* We first write down the explicit form of an arbitrary entry $CAM[i, j]$:

$$\begin{aligned} CAM[i,j] &= \sum_k (AM+I)[i,k] \cdot (AM+I)[j,k] \\ &= \sum_k AM[i,k]AM[j,k] + AM[i,k]I[j,k] + AM[j,k]I[i,k] + I[i,k]I[j,k] \\ &= \sum_k AM[i,k]AM[j,k] + \sum_k AM[i,k]I[j,k] + \sum_k AM[j,k]I[i,k] + \sum_k I[i,k]I[j,k] \end{aligned}$$

Since $AM$ is non-negative, we conclude that $\boldsymbol{CAM}[i, j] > 0$ if and only there exists at least one index $k$ such that at least one of the following conditions hold:

1. $\boldsymbol{AM}[i, k]\boldsymbol{AM}[j, k] > 0$
2. $\boldsymbol{AM}[i, k]\boldsymbol{I}[j, k] > 0$
3. $\boldsymbol{AM}[j, k]\boldsymbol{I}[i, k] > 0$

4. $\boldsymbol{I}[i,k]\boldsymbol{I}[j,k] > 0$

If condition 1 holds, individual $k$ is a common ancestor of individuals $i$ and $j$. If condition 2 holds, $j = k$ and individual $j$ is an ancestor of individual $i$. If condition 3 holds, the converse statement holds. Finally, if condition 4 holds, $i = j = k$, and so the individuals trivially share an ancestor.

■ □

##### 3.1.2 Creating the IBD matrix

After removing uninformative individuals, we compute the IBD matrix for the remaining individuals via dynamic programming. The algorithm we used is an efficient implementation of Lou's algorithm [15] for creating an IBD matrix from pedigree records, using Henderson's decomposition [7], and is detailed in Algorithm 1.

**Algorithm 1** Computing an IBD matrix

**Definitions**

- $CSRM(n,m)$ - Compressed Sparse Row $\mathbb{R}^{n\times m}$ matrix [17]
- $RLL\_SM(n,m)$ - Row-based linked list sparse $\mathbb{R}^{n\times m}$ matrix

**Paramethers**

- $ParentsList$ - individuals' parents

**Algorithm**

$\boldsymbol{L = RLL\_SM(N,N)}$
$H,\ F = \bar{0}_N$
**for** $i = 0;\ \ i < n$ **do**
    $L[i,i] = 1;\ \ ANC = i;\ \ iParents = ParentsList[i]$
    $H[i] = 1 - 0.25 \cdot (count(iParents) + sum(F[iParents]))$
    **while** $count(ANC) > 0$ **do**
        $j = max(ANC);\ jParents = AncestorList[j]$
        $L[i, jParents] \mathrel{+}= 0.5 \cdot L[i,j]$
        $F[i] \mathrel{+}= L[i,j]^2 \cdot H[i]$
        $ANC = ANC\ \cup jParents \setminus \{j\}$
    **end while**
    $F[i] \mathrel{-}= 1$
**end for**
$L\_CSRM = CSRM(L)$
$H\_CSRM = CSRM(withdiagonalH)$
**return** $L\_CSRM \times H\_CSRM \times L\_CSRM^T$

#### 3.2 Dominance matrix

The dominance kinship coefficient between individuals $i$ and $j$ is given by:

$$D_{i,j} = A_{p1_i,p2_j} \cdot A_{p2_i,p1_j} + A_{p1_i,p1_j} \cdot A_{p2_i,p2_j}. \tag{3}$$

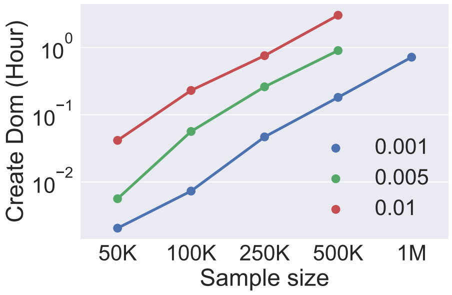

(a) Runtime of dominance matrix creation

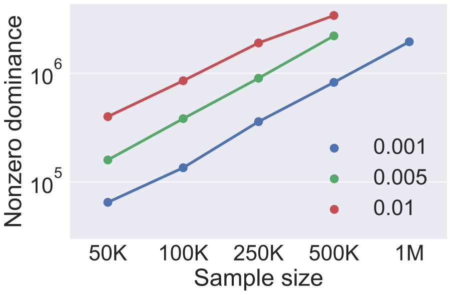

(b) #non-zero entries in the dominance matrix

Supplementary Figure 2: Average creation runtime and sparsity of simulated dominance matrices, under various sparsity factors.

Here, $A_{ij}$ is the IBD kinship coefficient of individuals $i$ and $j$, $p_{1_i}$, $p_{2_i}$ are the two parents of individual $i$. It is easy to compute the dominance matrix given an IBD matrix, because if $A_{ij} = 0$ then we also have $D_{ij} = 0$ (Lemma 2). Hence, we can compute the dominance matrix efficiently by only iterating over pairs of individuals with $A_{ij} = 0$ and computing the corresponding $D_{ij}$ entries (Supplementary Figure 2).

**Lemma 2.** *$A_{i,j} = 0$ if and only if $D_{i,j} = 0$.*

*Proof.* Let $i \neq j \in [N]$ such that $A_{i,j} = 0$. $\forall v \in [N]$ denote $f_v$ and $m_v$ as the different parents of node v. Suppose that $D_{i,j} \neq 0$. Therefore

$$A_{f_i,m_j} \cdot A_{m_i,f_j} + A_{f_i,f_j} \cdot A_{m_i,m_j} \neq 0$$

$$\forall k,l \in [N] A_{k,l} \geq 0 \Rightarrow \left(A_{f_i,m_j} \cdot A_{m_i,f_j} > 0\right) \text{ or } \left(A_{f_i,f_j} \cdot A_{m_i,m_j}\right) > 0$$

without loss of generality,

$$A_{f_i,m_j} \cdot A_{m_i,f_j} > 0 \Rightarrow A_{f_i,m_j} > 0$$

Therefore, given the definition of the IBD matrix, we are given an ancestor $o$ of both $f_i$ and $m_i$, making it a common ancestor of both $i$ and $j$, resulting in a non-zero $A_{i,j}$, contradicting our assumptions.

■ □

#### 4 Real Data Analysis

Our analysis of the GENI dataset closely followed the one described in [10]. Briefly, we first pruned non-informative individuals, as described in [10]. Afterwards, we applied a second round of pruning of uninformative individuals, as described in Section 3.1.1. This pruning stage reduced the cohort size from $43 \times 10^6$ to $1.6 \times 10^6$.

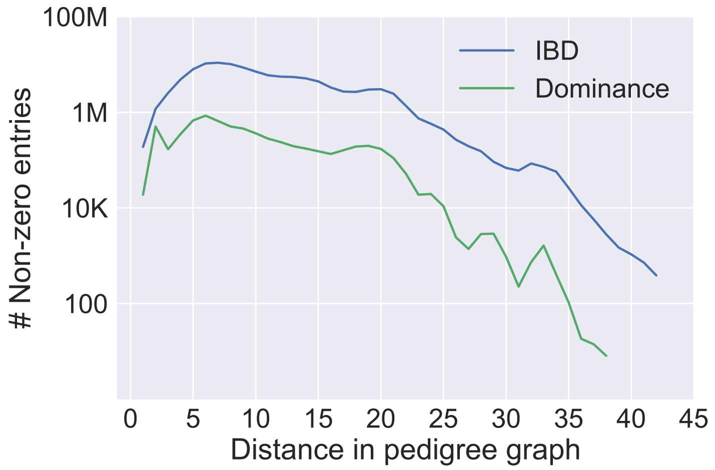

Supplementary Figure 3: The number of nonzero IBD and Dominance entries in the GENI dataset, as a function of degree of relationship

Afterwards, we computed an IBD matrix as described in Section 3.1, and a pairwise epistatis matrix, which consisted of the squares of the entries in the IBD matrix.

We did not include a dominance matrix in the estimation procedure, because it was extremely similar to the identity matrix. This is because of the relatively small number of parent-child links where both the parents and the children have full information (i.e., year of birth, year of death, and gender; Supplementary Figure 3). The sum of the squares of all entries in the dominance matrix was almost the same as the sum of the squares of the diagonal entries only (up to 0.001% error). We therefore opted to not include this matrix in the analysis.

### 5 Haseman Elston Estimator

Recall from the main text that the LMM Haseman Elston (HE) estimator requires fitting the following moment estimator:

$$E\left[\left(y_i - \boldsymbol{C}_i\boldsymbol{\beta}\right)\left(y_j - \boldsymbol{C}_j\boldsymbol{\beta}\right)\right] = \sum_{k=1}^{d} \sigma_k^2 M_{ij}^k, \tag{4}$$

where $y_i$,$y_j$ and $\boldsymbol{C}_i$, $\boldsymbol{C}_j$ are the phenotypes and covariate vectors of individuals $i$ and $j$, respectively, $\boldsymbol{\beta}$ is a vector of fixed effects, $d$ is the number of variance components, and $M_{ij}^k$ is the entry for individuals $i, j$ in variance component $k$.

The HE estimator can be evaluated in two steps. First, we estimate the fixed effects $\boldsymbol{\beta}$ via a standard linear regression model:

$$y_i = \boldsymbol{C}_i\boldsymbol{\beta} + \epsilon_i \tag{5}$$

Afterwards we plug the fixed effect estimate $\hat{\boldsymbol{\beta}}$ into Equation 4.

In the second step, we compute the variance component estimates $\hat{\sigma}_1^2, ..., \hat{\sigma}_d^2$ as follows:

$$\left[\hat{\sigma}_1^2, ..., \hat{\sigma}_d^2\right]^T = \left(\left[\boldsymbol{V}^1, ..., \boldsymbol{V}^d\right]^T \left[\boldsymbol{V}^1, ..., \boldsymbol{V}^d\right]\right)^{-1} \left[\boldsymbol{V}^1, ..., \boldsymbol{V}^d\right]^T \boldsymbol{Y}, \tag{6}$$

where $\boldsymbol{V}^k$ is a vector representation enumerating the elements $M_{ij}^k$ for all pairs of distinct individuals $i, j$, and $\boldsymbol{Y}$ is a vector representation of the corresponding elements $\left(y_i - \boldsymbol{C}_i\hat{\boldsymbol{\beta}}\right)\left(y_j - \boldsymbol{C}_j\hat{\boldsymbol{\beta}}\right)$. Each element $q, r$ of the $d \times d$ matrix $\left(\left[\boldsymbol{V}^1, ..., \boldsymbol{V}^d\right]^T \left[\boldsymbol{V}^1, ..., \boldsymbol{V}^d\right]\right)$ can be computed via an element-wise multiplication of the upper-diagonal elements of the matrices $\boldsymbol{M}^q$, $\boldsymbol{M}^r$, which can be performed efficiently via sparse matrix routines. The vector $\left[\boldsymbol{V}^1, ..., \boldsymbol{V}^d\right]^T \boldsymbol{Y}$ can also be computed efficiently in a similar manner.

Specifically, denoting $\mathbf{q} \triangleq \left[\boldsymbol{V}^1, ..., \boldsymbol{V}^d\right]^T \boldsymbol{Y}$, $\mathbf{S} = \left[\boldsymbol{V}^1, ..., \boldsymbol{V}^d\right]^T \left[\boldsymbol{V}^1, ..., \boldsymbol{V}^d\right]$, we have:

$$\left[\hat{\sigma}_1^2, ..., \hat{\sigma}_d^2\right]^T = \mathbf{S}^{-1}\mathbf{q}. \tag{7}$$

By applying a few matrix manipulations, we can compute $\mathbf{q}$ and $\mathbf{S}$ efficiently as follows:

$$\begin{aligned} q_k &= \mathbf{y}^T\mathbf{M}^k\mathbf{y} - \sum_i M_{ii}^k y_i^2 = \mathbf{y}^T\left(\mathbf{M}^k - \mathbf{I}\right)\mathbf{y} \\ \mathbf{S}_{kl} &= \sum_{ij} M_{ij}^k M_{ij}^l - \sum_i \mathbf{M}_{ii}^k \mathbf{M}_{ii}^l, \end{aligned} \tag{8}$$

where we used the assumption $M_{ii}^k = 1$. Both these quantities can be computed explicitly via sparse matrix routines.

The sampling variance of the estimators is given by $\mathbf{S}^{-1}\text{var}(\mathbf{q})\mathbf{S}^{-1}$, where $\text{var}(\mathbf{q})$ is given by:

$$\text{var}(\mathbf{q})_{kl} = 2\text{tr}\left(\mathbf{H}\left(\mathbf{M}^k - \mathbf{I}\right)\mathbf{H}\left(\mathbf{M}^l - \mathbf{I}\right)\right), \tag{9}$$

and where $\mathbf{H} = \sum_k \hat{\sigma}_k^2 \mathbf{M}^k$ is the covariance matrix of $\mathbf{y}$ [20]. This quantity can be computed in one of two ways:

1. Exactly, via: $\text{tr}\left(\mathbf{H}\left(\mathbf{M}^k - \mathbf{I}\right)\mathbf{H}\left(\mathbf{M}^l - \mathbf{I}\right)\right) = \sum_{ij}\left[\mathbf{H}\left(\mathbf{M}^k - \mathbf{I}\right)\right]_{ij}\left[\mathbf{H}\left(\mathbf{M}^l - \mathbf{I}\right)\right]_{ij}$
2. Approximately, via:
$$\text{tr}\left(\mathbf{H}\left(\mathbf{M}^k - \mathbf{I}\right)\mathbf{H}\left(\mathbf{M}^l - \mathbf{I}\right)\right) = \text{E}_{\mathbf{y}'}\left[\mathbf{y}'^T\mathbf{H}\left(\mathbf{M}^k - \mathbf{I}\right)\mathbf{H}\left(\mathbf{M}^l - \mathbf{I}\right)\mathbf{y}'\right], \tag{10}$$
where $\mathbf{y}'$ is sampled from $\mathcal{N}(\mathbf{0}, \mathbf{I})$.

The approximate approach uses Monte-Carlo approximations, analogously to the derivation of Equation 16. It can be substantially faster than the exact approach (because it can circumvent expensive matrix-matrix multiplications) and obtain almost the same accuracy, by sampling ~100 $\mathbf{y}'$ vectors. .

#### 6 Restricted Maximum Likelihood Estimator

Here we describe the efficient REML estimator for sparse LMMs. We first describe maximum likelihood (ML) estimation, and then describe how to extend the solution to REML. The solution presented here applied to general sparse LMMs. A more efficient solution for LMMs with a single variance component is presented in the next section.

##### 6.1 Maximum Likelihood Estimation

Recall that an LMM for a sample of $n$ individuals is defined as follows:

$$\boldsymbol{y} \sim \mathcal{N}(\boldsymbol{C\beta}\,;\ \sum_{k=1}^{d} \sigma_k^2 \boldsymbol{M}^k + \sigma_e^2 \boldsymbol{I}), \tag{11}$$

where $\boldsymbol{I}$ is the $n \times n$ identity matrix. We are interested in finding the maximum likelihood estimates (MLEs) of $\boldsymbol{\beta}$ and of $\sigma_1^2, ..., \sigma_d^2$, $\sigma_e^2$.

The LMM log likelihood is explicitly given by:

$$\ell(\boldsymbol{\beta}, \sigma_1^2, ..., \sigma_d^2, \sigma_e^2) = -\frac{1}{2}(\boldsymbol{y} - \boldsymbol{C\beta})\boldsymbol{K}^{-1}(\boldsymbol{y} - \boldsymbol{C\beta}) - \frac{1}{2}\log|\boldsymbol{K}| - \frac{n}{2}\log(2\pi), \tag{12}$$

where $\boldsymbol{K} = \sum_{k=1}^{d} \sigma_k^2 \boldsymbol{M}^k + \sigma_e^2 \boldsymbol{I}$ is the overall covariance matrix. To compute Equation 12 we need to compute the terms $\boldsymbol{K}^{-1}(\boldsymbol{y} - \boldsymbol{C\beta})$ and $\log|\boldsymbol{K}|$. The first term can be computed exactly via either conjugate gradient iterations [5] or by explicitly computing the Cholesky factorization of $\boldsymbol{K}$ and then applying forward and back substitution [5]. The second term can be computed via the Cholesky factorization of $\boldsymbol{K}$. The Cholesky factorization can be computed efficiently via the CHOLMOD routines [3]. It remains to find the maximum likelihood estimates of the model parameters.

To find the MLE of $\hat{\boldsymbol{\beta}}$ we note that given $\boldsymbol{K}$, $\hat{\boldsymbol{\beta}}$ can be computed analytically by deriving Equation 12 with respect to $\boldsymbol{\beta}$ as follows:

$$\frac{\partial \ell(\boldsymbol{\beta}, \sigma_1^2, ..., \sigma_d^2, \sigma_e^2)}{\partial \boldsymbol{\beta}} = -\frac{1}{2}(y - \boldsymbol{C\beta})^T \boldsymbol{K}^{-1}\boldsymbol{C} \tag{13}$$

By setting the transpose of the gradient to 0, we obtain the MLE:

$$\hat{\boldsymbol{\beta}} = \left(\boldsymbol{C}^T \boldsymbol{K}^{-1} \boldsymbol{C}\right)^{-1} \boldsymbol{C}^T \boldsymbol{K}^{-1} \boldsymbol{y}. \tag{14}$$

The MLEs of the variance components $\hat{\sigma}_1^2, ..., \hat{\sigma}_d^2$ are estimated via an optimization procedure, which requires computing the gradient of Equation 12. The partial derivative with respect to each variance components $\sigma_k^2$ is given by:

$$\frac{\partial \ell(\boldsymbol{\beta}, \sigma_1^2, ..., \sigma_d^2, \sigma_e^2)}{\partial \sigma_k^2} = -\frac{1}{2}\boldsymbol{y}^T \boldsymbol{K}^{-1} \boldsymbol{M}^k \boldsymbol{K}^{-1} \boldsymbol{y} - \frac{1}{2}\operatorname{Tr}\left[\boldsymbol{K}^{-1}\boldsymbol{M}^k\right]. \tag{15}$$

The first term on the right hand side of Equation 15 can be computed efficiently given the Cholesky factorization of $\boldsymbol{K}$. Unfortunately, the second term cannot be solved efficiently via the above technique because it requires solving $n$ different linear equations, where $n$ can be in the millions. Instead, we use the approximation technique proposed in [14]. We first rewrite this term as an expectation (ignoring the scaling factor) as follows:

$$\begin{aligned}
\operatorname{Tr}\left[\boldsymbol{K}^{-1}\boldsymbol{M}^k\right] &= \operatorname{Tr}\left[\boldsymbol{K}^{-1}\boldsymbol{M}^k\boldsymbol{K}^{-1}\boldsymbol{K}\right] \\
&= \operatorname{Tr}\left[\boldsymbol{K}^{-1}\boldsymbol{M}^k\boldsymbol{K}^{-1}E\left[\boldsymbol{y}'\boldsymbol{y}'^T\right]\right] \\
&= E\left[\operatorname{Tr}\left[\boldsymbol{K}^{-1}\boldsymbol{M}^k\boldsymbol{K}^{-1}\boldsymbol{y}'\boldsymbol{y}'^T\right]\right] \\
&= E\left[\operatorname{Tr}\left[\boldsymbol{y}'^T\boldsymbol{K}^{-1}\boldsymbol{M}^k\boldsymbol{K}^{-1}\boldsymbol{y}'\right]\right] \\
&= E\left[\boldsymbol{y}'^T\boldsymbol{K}^{-1}\boldsymbol{M}^k\boldsymbol{K}^{-1}\boldsymbol{y}'\right],
\end{aligned} \tag{16}$$

where $\boldsymbol{y}' \sim \mathcal{N}(\mathbf{0}, \boldsymbol{K})$ and we used the fact that the trace of a scalar is equal to the scalar. We therefore approximate Equation 16 by sampling a small number of $\boldsymbol{y}'$ vectors to approximate the expectation. These vectors can be sampled efficiently given the Cholesky factorization $\boldsymbol{K} = \boldsymbol{L}\boldsymbol{L}^T$ by sampling a vector $\boldsymbol{y}_K$ of $n$ independent variables sampled from $\mathcal{N}(0,1)$ and then using the fact that $\boldsymbol{L}\boldsymbol{y}_K \sim \mathcal{N}(\mathbf{0}, \boldsymbol{K})$. The Cholesky factorization can be computed efficiently via the CHOLMOD routines. We found that 100 vectors often yields a very good approximation at a negligible computational cost.

We note that [14] proposes an alternative estimation method by completely foregoing the likelihood computation, and instead only trying to minimize the squared gradient elements. However, we found that in sparse settings, this solution often converges into local maxima at the edge of the parameter space (where many variance components are equal to zero) rather than the true maximum likelihood estimate.

##### 6.2 Restricted Maximum Likelihood Estimation

The restricted log likelihood is given by:

$$\ell_R(\boldsymbol{\beta}, \sigma_1^2, ..., \sigma_d^2, \sigma_e^2) = \ell(\boldsymbol{\beta}, \sigma_1^2, ..., \sigma_d^2, \sigma_e^2) + \frac{c}{2}\log(2\pi) + \frac{1}{2}\log\left|\boldsymbol{C}^T\boldsymbol{C}\right| - \frac{1}{2}\log\left|\boldsymbol{C}^T\boldsymbol{K}^{-1}\boldsymbol{C}\right|, \tag{17}$$

where $c$ is the number of covariates. This expression can be shown to be equivalent to the expression presented in the main text [13].

Clearly, the restricted maximum likelihood estimate (RMLE) of $\boldsymbol{\beta}$ is the same as the MLE. The derivative of the restricted log likelihood with respect to each variance component $\sigma_k^2$ is given by:

$$\frac{\partial \ell_R(\boldsymbol{\beta}, \sigma_1^2, ..., \sigma_d^2, \sigma_e^2)}{\sigma_k^2} = \frac{\partial \ell(\boldsymbol{\beta}, \sigma_1^2, ..., \sigma_d^2, \sigma_e^2)}{\sigma_k^2} + \frac{1}{2}\operatorname{Tr}\left[\left(\boldsymbol{C}^T\boldsymbol{K}^{-1}\boldsymbol{C}\right)^{-1}\boldsymbol{C}^T\boldsymbol{K}^{-1}\boldsymbol{M}^k\boldsymbol{K}^{-1}\boldsymbol{C}\right]. \tag{18}$$

The term $\boldsymbol{K}^{-1}\boldsymbol{C}$ can be computed by solving $c$ different linear equations, which can be performed efficiently given the Cholesky factorization of $\boldsymbol{K}$. All the other terms can be computed efficiently, assuming that $c$ is small compared to $n$.

##### 6.3 Implementation details

We implemented the above optimization procedure in Python, using an L-BFGS-B algorithm [2] as implemented in the SciPy package [9]. To prevent the parameters from inducing a non positive-definite matrix, We enforced non-negative parameters by using a log-transformation, which transforms the problem into an unconstrained optimization problem.

#### 7 Fitting Sparse LMMs with a Single Variance Component

In the special case of a single variance component ($d$=1), we can use special techniques similar to those proposed in [12] to obtain an even more efficient solution. We now describe the estimation procedure for maximum likelihood and for restricted maximum likelihood estimates.

##### 7.1 Maximum Likelihood Estimation for a Single Variance Component

Denote $\boldsymbol{M}$ as the matrix associated with the variance component, and $\boldsymbol{K} = \sigma_g^2 \boldsymbol{M} + \sigma_e^2 \boldsymbol{I}$ as the covariance matrix of $\boldsymbol{y}$. To solve the problem efficiently, we introduce the variances ratio $\delta = \frac{\sigma_e^2}{\sigma_g^2}$ and define the matrix $\boldsymbol{U} = \boldsymbol{M} + \delta \boldsymbol{I}$. Note that $\boldsymbol{K} = \sigma_g^2 \boldsymbol{U}$. We can therefore reparameterize the distribution of $\boldsymbol{y}$ as follows:

$$\boldsymbol{y} \sim \mathcal{N}(\boldsymbol{C\beta}\,;\, \sigma_g^2 \boldsymbol{U}). \tag{19}$$

It is convenient to use this parameterization because the MLE of all the parameters can be found via closed-form formulas given $\delta$. We can therefore convert the problem from a $c+2$ dimensional problem into an effectively one dimensional problem, where we search for the value of $\delta$ that maximizes the likelihood.

We now describe how the MLEs of $\boldsymbol{\beta}$ and of the variance parameters can be computed given $\delta$. Similarly to the general LMM solution, the fixed effect estimates are given by:

$$\hat{\boldsymbol{\beta}} = \left(\boldsymbol{C}^T \boldsymbol{U}^{-1} \boldsymbol{C}\right)^{-1} \boldsymbol{C}^T \boldsymbol{U}^{-1} \boldsymbol{y}. \tag{20}$$

Note that $\hat{\boldsymbol{\beta}}$ is independent of $\sigma_g^2$ given $\delta$. We can therefore define $\tilde{\boldsymbol{y}} = \boldsymbol{y} - \boldsymbol{C}\hat{\boldsymbol{\beta}}$ as the value of $\boldsymbol{y}$ after regressing out the fixed effects.

The MLE of $\sigma_g^2$ is found by maximizing the profile likelihood:

$$\ell(\sigma_g^2) = -\frac{1}{2} \tilde{\boldsymbol{y}} \boldsymbol{K}^{-1} \tilde{\boldsymbol{y}} - \frac{1}{2} \log |\boldsymbol{K}| - \frac{n}{2} \log(2\pi). \tag{21}$$

To find the MLE, we derive Equation 21 according to $\sigma_g^2$. Using the formula for the partial derivative of a multivariate normal distribution, we obtain:

$$\begin{aligned} \frac{\partial \ell(\sigma_g^2)}{\partial \sigma_g^2} &= \frac{1}{2} \tilde{\boldsymbol{y}}^T \boldsymbol{K}^{-1} \frac{\partial \boldsymbol{K}}{\partial \sigma_g^2} \boldsymbol{K}^{-1} \tilde{\boldsymbol{y}} \,-\, \frac{1}{2} \mathrm{tr} \left( \boldsymbol{K}^{-1} \frac{\partial \boldsymbol{K}}{\partial \sigma_g^2} \right) \\ &= \frac{1}{2} \tilde{\boldsymbol{y}}^T \left( \sigma_g^{-2} \boldsymbol{U}^{-1} \boldsymbol{M} \sigma_g^{-2} \boldsymbol{U}^{-1} \right) \tilde{\boldsymbol{y}} \,-\, \frac{1}{2} \sigma_g^{-2} \,\mathrm{Tr}\left[ \boldsymbol{U}^{-1} \boldsymbol{M} \right]. \end{aligned} \tag{22}$$

By setting the partial derivative to 0, we obtain the MLE:

$$\hat{\sigma}_g^2 = \frac{\tilde{\boldsymbol{y}}^T \boldsymbol{U}^{-1} \boldsymbol{M} \boldsymbol{U}^{-1} \tilde{\boldsymbol{y}}}{\mathrm{Tr}\left[\boldsymbol{U}^{-1} \boldsymbol{M}\right]} = \frac{\left(\boldsymbol{U}^{-1} \tilde{\boldsymbol{y}}\right)^T \boldsymbol{M} \left(\boldsymbol{U}^{-1} \tilde{\boldsymbol{y}}\right)}{\mathrm{Tr}\left[\boldsymbol{U}^{-1} \boldsymbol{M}\right]} \tag{23}$$

As in the general case, the numerator can be solved efficiently using either conjugate gradient iterations or the Cholesky factorization of $\boldsymbol{U}$, and the denominator can be accurately approximated via an expectation. Specifically, we apply the following approximation:

$$\mathrm{Tr}\left[\boldsymbol{U}^{-1} \boldsymbol{M}\right] = E_{\boldsymbol{y}_U} \left[ \boldsymbol{y}_U^T \boldsymbol{U}^{-1} \boldsymbol{M} \boldsymbol{U}^{-1} \boldsymbol{y}_U \right], \tag{24}$$

where $\boldsymbol{y}_U$ is a random vector with the distribution $\boldsymbol{y}_U \sim \mathcal{N}(\mathbf{0}\,; \boldsymbol{U})$, and the expectation is approximated via 100 randomly sampled $\boldsymbol{y}_U$ vectors. Drawing random vectors $\boldsymbol{y}_U$ is easy when we have the Cholesky decomposition of $\boldsymbol{M}$. Specifically, denoting the Cholesky decomposition via $\boldsymbol{M} = \boldsymbol{L}\boldsymbol{L}^T$ (where $\boldsymbol{L}$ is a lower triangular matrix) and recalling that $\boldsymbol{U} = \boldsymbol{M} + \delta\boldsymbol{I}$, we can draw $\boldsymbol{y}_U$ vectors by first sampling two vectors of independent standard normal variables $\boldsymbol{z}_L$, $\boldsymbol{z}_I$ and then sampling $\boldsymbol{y}_U$ via $\boldsymbol{y}_U = \boldsymbol{L}\boldsymbol{z}_L + \delta\boldsymbol{z}_I$. Note that technically we could use the fact that we have the factorization of $\boldsymbol{U}$ and use its own Cholesky decomposition, but this is more computationally expensive because we have to explicitly compute the lower diagonal matrix for each evaluated value of $\delta$, instead of using the same matrix $\boldsymbol{L}$ for all values of $\delta$.

Finally, we need to estimate the MLE of $\sigma_e^2$, but it is trivially given via the relation $\sigma_e^2 = \delta\sigma_g^2$.

After computing the Cholesky decomposition of $\boldsymbol{U}$ and obtaining the MLE of $\boldsymbol{\beta}$, $\sigma_g^2$ and $\sigma_e^2$ for a specific value of $\delta$, we can compute the log likelihood (Equation 12) via:

$$\begin{aligned}\ell(\boldsymbol{\beta}, \sigma_g^2, \delta) &= -\frac{1}{2}(y - \boldsymbol{C}\boldsymbol{\beta})\left(\sigma_g^2\boldsymbol{U}\right)^{-1}(y - \boldsymbol{C}\boldsymbol{\beta}) - \frac{1}{2}\log\left|\sigma_g^2\boldsymbol{U}\right| - \frac{n}{2}\log(2\pi) \\ &= -\frac{1}{2}\sigma_g^{-2}\tilde{\boldsymbol{y}}^T\boldsymbol{U}^{-1}\tilde{\boldsymbol{y}} - \frac{n}{2}\log\sigma_g^2 - \frac{1}{2}\log|\boldsymbol{U}| - \frac{n}{2}\log(2\pi). \end{aligned} \tag{25}$$

The product $\boldsymbol{U}^{-1}\tilde{\boldsymbol{y}}$ and and the log determinant $\log|\boldsymbol{U}|$ can easily be computed given the Choleksy decomposition of $\boldsymbol{U}$.

The optimization procedure to estimate the MLEs is carried out as follows. We iterate over a grid of $\log\delta$ values in the range $[-5, 5]$ (this should cover the entire spectrum for both very small and very large ratios of $\sigma_e^2$ to $\sigma_g^2$). For each evaluated value of $\delta$, we first compute the Cholesky decomposition of $\boldsymbol{U} = \boldsymbol{M} + \delta\boldsymbol{I}$. Using this decomposition we can first estimate the fixed effects $\boldsymbol{\beta}$ via Equation 20, then we estimate $\sigma_g^2$ via Equation 23 and finally we estimate $\sigma_e^2$ via $\hat{\sigma}_e^2 = \delta\hat{\sigma}_g^2$. The selected value is the one that maximizes the log likelihood, as computed in Equation 25.

#### 7.2 Restricted Maximum Likelihood Estimation for a Single Variance Component

Recall that the restricted log likelihood is given by:

$$\ell_R(\boldsymbol{\beta}, \sigma_g^2, \delta) = \ell(\boldsymbol{\beta}, \sigma_g^2, \delta) + \frac{c}{2}\log(2\pi) + \frac{1}{2}\log\left|\boldsymbol{C}^T\boldsymbol{C}\right| - \frac{1}{2}\log\left|\boldsymbol{C}^T\boldsymbol{V}^{-1}\boldsymbol{C}\right|. \tag{26}$$

Clearly, the restricted maximum likelihood estimate (RMLE) of $\boldsymbol{\beta}$ is the same as the MLE. The derivative of $\ell_R(\sigma_g^2)$ with respect to $\sigma_g^2$ is given by:

$$\begin{aligned}\frac{\partial\ell_R(\sigma_g^2)}{\partial\sigma_g^2} &= \frac{\partial\ell(\sigma_g^2)}{\partial\sigma_g^2} + \frac{1}{2}\mathrm{tr}\left(\left(\boldsymbol{C}^T\boldsymbol{K}^{-1}\boldsymbol{C}\right)^{-1}\boldsymbol{C}^T\boldsymbol{K}^{-1}\frac{\partial\boldsymbol{K}}{\partial\sigma_g^2}\boldsymbol{V}^{-1}\boldsymbol{C}\right) \\ &= \frac{\partial\ell(\sigma_g^2)}{\partial\sigma_g^2} + \frac{1}{2}\sigma_g^{-2}\mathrm{tr}\left(\left(\boldsymbol{C}^T\boldsymbol{U}^{-1}\boldsymbol{C}\right)^{-1}\boldsymbol{C}^T\boldsymbol{U}^{-1}\boldsymbol{M}\boldsymbol{U}^{-1}\boldsymbol{C}\right). \end{aligned} \tag{27}$$

By combining Equations 27 and 22, the RMLE of $\sigma_g^2$ is given by:

$$\hat{\sigma}_g^2 = \frac{\left(\boldsymbol{U}^{-1}\tilde{\boldsymbol{y}}\right)^T \boldsymbol{M} \left(\boldsymbol{U}^{-1}\tilde{\boldsymbol{y}}\right)}{\text{tr}\left(\boldsymbol{U}^{-1}\boldsymbol{M}\right) + \text{tr}\left(\left(\boldsymbol{C}^T\boldsymbol{U}^{-1}\boldsymbol{C}\right)^{-1}\boldsymbol{C}^T\boldsymbol{U}^{-1}\boldsymbol{M}\boldsymbol{U}^{-1}\boldsymbol{C}\right)}. \tag{28}$$

All the terms in the second term of the denominator are easy to compute, assuming that the number of covariates $c$ is very small compared to $n$. We can therefore repeat the procedure described in the previous section for REML estimation.

### 8 Estimating Standard Errors of Variance Component Estimates

We estimated the standard errors of the variance component estimates via the average information REML (AI-REML) procedure [4], which consists of approximating the Hessian of the log likelihood as follows:

$$\frac{\partial \ell(\boldsymbol{\beta}, \sigma_1^2, ..., \sigma_d^2, \sigma_e^2)}{\sigma_k^2 \sigma_l^2} \approx -\frac{1}{2}\boldsymbol{y}^T\boldsymbol{K}^{-1}\boldsymbol{M}^k\boldsymbol{K}^{-1}\boldsymbol{M}^l\boldsymbol{K}^{-1}\boldsymbol{y}. \tag{29}$$

It is straightforward to compute Equation 29 with sparse matrices, given their Cholesky decomposition.

We note that when using REML estimation, the matrix $\boldsymbol{K}^{-1}$ in Equation 29 is replaced by $\tilde{\boldsymbol{K}} \triangleq \boldsymbol{K}^{-1} - \boldsymbol{K}^{-1}\boldsymbol{C}\left(\boldsymbol{C}^T\boldsymbol{K}^{-1}\boldsymbol{C}\right)^{-1}\boldsymbol{C}^T\boldsymbol{K}^{-1}$.

[10] Joanna Kaplanis, Assaf Gordon, Tal Shor, Omer Weissbrod, Dan Geiger, Mary Wahl, Michael Gershovits, Barak Markus, Mona Sheikh, Melissa Gymrek, Gaurav Bhatia, Daniel G MarArthur, Alkes Price, and others. Quantitative analysis of population-scale family trees using millions of relatives. *Science, in press*, page 106427, 2017.

[18] U.S. Census Bureau. Number, timing, and duration of marriages and divorces: 2009. U.S. Department of Commerce, may 2011.

[19] U.S. Census Bureau. America's families and living arrangements: 2012. U.S. Department of Commerce, august 2013.

[20] Xiang Zhou. A unified framework for variance component estimation with summary statistics in genome-wide association studies. *The annals of applied statistics*, 11(4):2027, 2017.
